# Preparatory encoding of diverse features of intended movement in the human motor cortex

**DOI:** 10.1101/2025.09.24.678356

**Authors:** Mattia Rigotti-Thompson, Samuel R. Nason-Tomaszewski, Payton Bechefsky, Alexander Acosta, Nick Hahn, Donald Avansino, Brice Richards, Claire Nicolas, Yahia H. Ali, Jaimie M. Henderson, Leigh R. Hochberg, Nicholas AuYong, Chethan Pandarinath

**Author notes:** These authors contributed equally (Nicholas AuYong, Chethan Pandarinath).

## Abstract

Over the course of a voluntary movement, motor cortical activity exhibits a transition from preparation to execution, with markedly different activity across these phases. Preparatory activity in particular might be used to improve brain-computer interfaces (BCIs) that harness brain activity to control external assistive devices, for example by anticipating a user’s intended movement trajectory for quick and fluid performance. However, to leverage preparatory activity for clinical BCIs, we must first understand which features of upcoming movements are encoded by preparatory activity in humans. In this work, we collected intracortical recordings from 3 research participants in the BrainGate2 clinical trial to investigate whether diverse features of movement, such as direction, curvature, and distance, are encoded by preparatory activity in the human motor cortex. We first show that preparatory activity is tuned to the direction of upcoming movements, and this tuning is largely preserved across movements with different effectors. Further investigation demonstrated this preparatory activity is also informative of initial and endpoint directions of curved movement trajectories, and encodes movement distance and speed independently. Finally, we present an online control paradigm that leverages preparatory activity to predict movements towards intended directions in advance, yielding rapid, self-paced control of a computer cursor by human participants. Altogether, these results demonstrate that preparatory activity in the human motor cortex encodes rich features of upcoming movement, highlighting its potential use for high performance brain-computer interface applications.

## Introduction

Voluntary movement execution is often preceded by a period of movement preparation^1^. This preparation process optimizes the performance of the upcoming movement for a particular goal, as evidenced by behavior becoming more precise with longer preparation times^2,3^. Neurophysiological studies support a neural basis for motor preparation, as spiking activity that precedes movement execution is an inherent property of neuron populations within the motor cortex^4–6^. This “preparatory” activity is distinct from the neural activity observed during movement execution and can arise several seconds before any actual motor output^7–9^. Interestingly, this preparatory activity also arises in the absence of any explicitly allotted preparation time^10–13^, highlighting it as a critical component of neural activity in the motor cortex related to movement generation. In this regard, its role is often linked to a priming mechanism that sets an optimal neural state for the upcoming movement^1,14,15^.

A considerable body of non-human primate research has centered on identifying which aspects of upcoming movement are encoded in preparatory activity. By using instructed-delay paradigms that explicitly separate preparatory and execution-related activity, studies with non-human primates have demonstrated that preparatory activity is informative of several properties of upcoming movement. These properties can be temporal, such as reaction time and movement speed^16–20^, or spatial such as movement direction, distance, and curvature^4,21–24^.

Given the apparent richness of preparatory activity, it might be used to improve brain-computer interfaces (BCIs) which aim to predict a user’s intended movement from neural recordings to control assistive devices^25–29^. However, standard BCIs do not capitalize on the wealth of information from preparatory activity, and instead rely on mapping execution-related neural activity onto instantaneous movement commands for an external effector (kinematics, force, etc.). Such paradigms can produce inaccurate movement trajectories due to the characteristic moment-to-moment variability in neural activity. An alternative BCI control paradigm might leverage preparatory information in neural activity to predict features of intended movement in advance. For example, a user’s intended movement trajectory could be precomputed during preparation to quickly and fluidly control movement of an external effector. Previous work in non-human primates has shown that, at least during simple motor tasks, BCIs that predict movement goals in advance can result in fast and accurate control^30–32^.

Nonetheless, few studies have examined preparatory activity from the motor cortex of humans, particularly from people who are unable to perform overt movements as is the usual case for implantable BCI users. In other brain areas associated with motor planning and sensory integration (such as posterior parietal cortex), activity that precedes movement is expectedly prominent, and tuning to some features of upcoming movement (e.g. direction or selected action) have been identified^33,34^. Within the human motor cortex itself, preparatory activity has also been shown to be distinct between simple movements, such as towards different target directions^35^, or involving different finger groupings^34^. However, BCI control paradigms built around precomputing intended movement trajectories in advance would require a thorough understanding of how movement features that encompass a broad repertoire of motor behaviors are represented in human preparatory activity. For example, movement curvature and distance are critical features for BCI control whose preparatory encoding has been underexplored in humans.

In this work, we studied the preparatory neural activity that precedes movement in the motor cortex of humans through intracortical recordings from three research participants enrolled in the BrainGate2 clinical trial. In all participants, the dorsal precentral gyrus exhibited preparatory activity that was strongly tuned to the direction of upcoming movement. A substantial component of this directional tuning was also preserved when attempting movements with different effectors. To characterize the tuning of preparatory activity to a richer assortment of movement features, we designed tasks that specifically probed movements with different curvatures, distances, and speeds. Through these tasks, we demonstrated that preparatory activity in human BCI participants with tetraplegia is also tuned to both the initial and endpoint directions of curved movement trajectories, and showed tuning to movement distance and speed independently of each other. Lastly, as a proof of concept for practical brain-computer interface use, we designed and tested an online control paradigm that leveraged preparatory activity to predict movements towards intended directions in advance, allowing rapid and self-paced control of an on-screen cursor for two BCI participants.

## Results

### The human motor cortex exhibits preparatory activity that is strongly tuned to upcoming movement direction

We collected multi-unit intracortical recordings from the hand-knob area of the precentral gyrus (putative Brodmann area 6d) from participants T16, T11, and T5 enrolled in the BrainGate2 clinical trial. To identify the presence of preparatory activity preceding movement initiation, participants performed an instructed-delay task using attempted movements of their wrists to match on-screen cursor movements (Fig. 1A). In each trial, one of eight possible targets (radial-8) was first highlighted (“target cue”; Fig. 1B). After a delay of random duration, participants were cued to attempt a quick movement towards the highlighted target (“go cue”). For T16 and T11, a random subset of trials skipped the go cue, such that the delay period was immediately followed by a rest period preceding the next trial. In these cases, participants were instructed not to attempt movements. For T5, a circular timer was shown during the delay period indicating the remaining delay time.

**Fig. 1:**
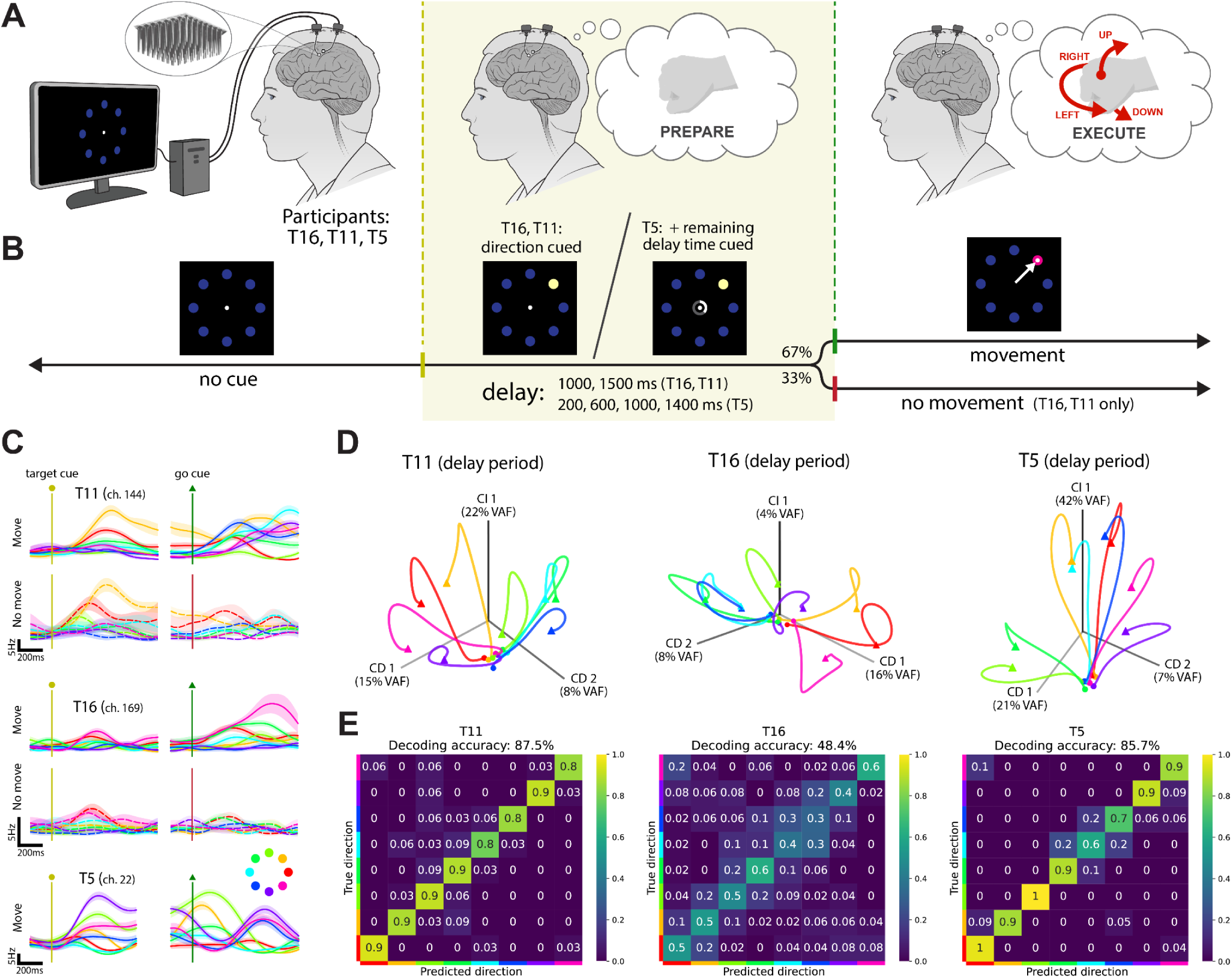
The hand-knob area of human participants exhibits preparatory activity that is strongly tuned to upcoming movement direction. **A.** Neural activity was recorded from intracortical microelectrodes in the hand-knob area of precentral gyrus in human participants preparing and attempting directional movements with their wrists. **B.** Trial structure and timing for the instructed-delay radial-8 task for the different participants with task visuals for an example trial. **C.** Average firing rates through time for example neural channels from each participant, for trials corresponding to the different movement directions and with a delay duration ≥ 1000 ms. For participants T11 and T16, solid lines correspond to trials where movement initiation was cued and dashed lines correspond to trials where they were instructed to skip movement. For all traces, the shaded regions indicate +/-the standard error of the mean. **D.** Average neural population activity during the delay period for each movement direction, as visualized by demixed principal component analysis. Projections onto the top condition-independent (“CI 1”) and condition-dependent (“CD 1” and “CD 2”) components are shown. Trajectories span the time from the target cue (indicated by a circle) to 1000 ms (T11, T16) or 800 ms (T5) after the target cue (indicated by a triangle). Trajectories are colored by movement direction, following the same color scheme as in C. **E.** Confusion matrices for single-trial movement direction predictions decoded from preparatory activity during the delay period, using support vector machine classifiers (leave-one-out cross-validation).

In all participants, neural channels exhibited sustained activity during the delay period that was tuned to the upcoming direction of movement and was distinct from execution period activity (Fig. 1C). This preparatory tuning was prominent: across the neural population, >50% of active channels from each participant were tuned to upcoming movement direction (*p*-value < 0.01, ANOVA; Supplemental Fig. 1). During trials when movement was not initiated, preparatory activity decayed back to a baseline state after the cue to cancel movement was shown, indicating that this activity is indeed tied to movement preparation and is independent from movement execution itself.

Changes in neural activity before the instructed execution period might relate to the participant attempting movements early rather than being tied to motor preparation. To rule out this possibility, we used surface electromyography (EMG) to record residual muscle activity from participant T16’s forearm while she performed the task. EMG modulation was only seen during the execution period, and was absent for trials where this period was skipped, confirming that movements were only attempted when instructed (Supplemental Fig. 2). Additionally, to test whether preparatory activity was due purely to visual stimuli from the cued targets, we designed an alternate version of the task aimed to reduce the targets’ visual saliency and resulting eye movements. In this version, movement direction was cued by an arrow at the center of the screen, and the participant was instructed to maintain their gaze on that center cue (Supplemental Fig. 3). Even with a lack of pronounced eye movements, preparatory activity was still present and consistent with that observed during the regular version of the task, confirming that some aspect of this preparatory activity is intrinsically tied to wrist movement intention.

We then visualized the dependence of preparatory activity on movement direction at the neural population level using demixed principal components analysis (dPCA)^36^. As shown, preparatory activity for each directional condition became distinct from baseline across time and exhibited structure across conditions that was consistent with the targets’ spatial distribution (Fig. 1D).

To evaluate how robust this preparatory representation of movement direction is on a single-trial basis, we trained support vector machine (SVM) classifiers to predict movement direction from neural population activity. Cross-validated decoding accuracies ranged from 48% to 88% (chance: 14%; Fig. 1E), which is noteworthy given that classification was restricted to only use neural activity prior to movement initiation. Taken together, these results confirm that preparatory activity is a characteristic feature of the neural activity in the human motor cortex with pronounced tuning to the direction of upcoming movement.

### Preparatory activity exhibits effector-independent directional tuning

While human motor cortex clearly exhibits prominent directional tuning in its preparatory activity, it is unclear if this tuning is tied to the specific effector used for movement (that is, the body part and associated muscles the participant attempts to move for the purposes of cursor control), or alternatively, tied to task-related movement intention in an effector-agnostic manner (e.g., “move right” or “move left”). Both tuning scenarios would be useful for BCI applications: the first would enable prediction of the body part a user plans to move, whereas the latter would enable robust direction decoding that generalizes across effectors. Recent work has shown that during movement execution, movements are represented in the hand-knob area of the human precentral gyrus in a compositional manner, with some components of neural activity tied to movement effector and other components preserved across analogous movements with different effectors^37^. However, it is uncertain if these findings hold for movement preparation, which is a distinct process that occurs before any effector-specific intended muscle output.

To evaluate the dependence of preparatory directional tuning on movement effector identity, we asked participants T16 and T11 to perform the radial-8 task using different effectors (Fig. 2A). Specifically, participants attempted movements of their fingers (small finger for T16, index finger for T11) or their wrists. For individual neural channels, preparatory activity showed a wide range of tuning variability between effectors: some channels had similar responses for wrist and finger movements, while others presented a pronounced change in directional tuning and/or modulation (Fig. 2B). At the population level, there was notable alignment between the activity corresponding to movements with different effectors but to the same direction, as indicated by positively correlated average delay period activity across effectors (mean correlations of 0.94 for T16 and 0.42 for T11 for off-diagonal values of pairwise neural correlation matrices; Fig. 2C). This demonstrates that some aspect of preparatory activity is effector-independent, that is, related to abstract task features rather than the specific attempted physical movement.

**Fig. 2:**
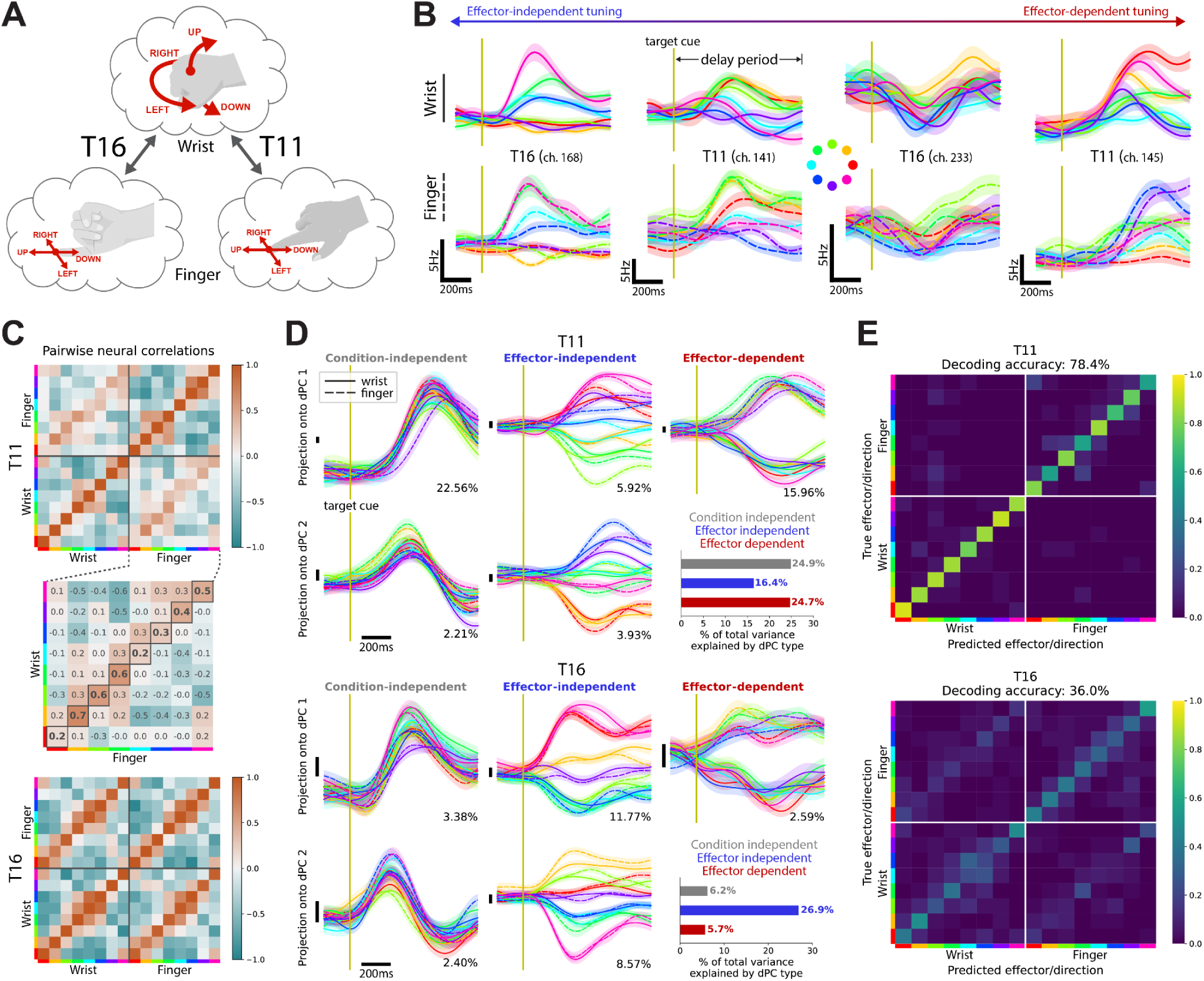
Preparatory activity exhibits effector-independent directional tuning. **A.** Participants were instructed to perform the radial-8 task (from Fig. 1) using attempted movements of their small finger (T16), index finger (T11), or wrist. **B.** Average firing rates through time for example neural channels during the delay period for trials using either the wrist (top row) or finger (bottom row). Shaded regions indicate +/-the standard error of the mean for each condition. **C.** Pairwise neural correlations between the delay period activity (averaged across time) across all movement conditions. Correlation was computed using a cross-validated estimator that reduces bias, and can result in values with magnitude greater than 1 (see Methods for details). **D.** Projection of average neural population activity during the delay period onto demixed principal components (dPCs) that are either condition-independent, effector-independent, or effector-dependent. Solid traces correspond to wrist movements and dashed traces correspond to finger movements. Traces are colored by movement direction (as indicated in B) and shaded regions indicate +/-the standard error of the mean for projections corresponding to trials from each condition. Vertical black bars to the left of each subplot indicate a consistent scale across dPC projections (arbitrary units). The number on the bottom right of each subplot indicates the percentage of total preparatory activity variance explained by each component. Bar plots on the bottom right show the percentage of the total trial-averaged preparatory activity variance accounted for by each dPC type. **E.** Confusion matrices for single-trial joint movement direction and effector predictions decoded from preparatory activity during the delay period, using SVM classifiers (leave-one-out cross-validation).

To quantify the degree of effector dependence, we used dPCA to linearly decompose preparatory activity into components that are either effector-dependent, effector-independent, or preserved across all conditions (effectors and movement directions). For both participants, dPCA revealed effector-independent components of preparatory activity that were consistently tuned to upcoming movement direction (Fig. 2D). There were also distinct effector-dependent components between movements with either the wrist or finger. The relative magnitudes of these components varied across participants: for T16, the effector-independent components accounted for a larger fraction of the total preparatory activity variance than the effector-dependent components, whereas the opposite was the case for participant T11.

Given the prominence of effector-independent information, it was unclear if both movement direction and effector identity could be jointly distinguished from preparatory activity. This is particularly important in practical brain-computer interface applications, where both pieces of information might be required. For this purpose, we evaluated how accurately both the direction and effector identity of upcoming movement could be predicted from preparatory activity on single trials, using SVM classifiers. Decoding accuracies ranged from 36% to 78% across participants, considerably above the chance level (chance: 7%; Fig. 2E). This suggests that preparatory activity is unique for each different combination of movement effector and direction, in line with theories stating its role is to set an initial neural state to drive the specific upcoming movement^14,15^.

### Preparatory activity is tuned to both the initial and endpoint directions of curved trajectories

In the previous sections, we established that preparatory activity was informative of the direction of upcoming movements. However, given that the attempted movements were all straight, it cannot be determined if the observed tuning was tied to the initial direction of movement or the direction of the trajectory endpoint (“target” direction).

To address this question, we asked participants T16 and T11 to perform a new task that instructed movements with a variety of straight and curved trajectories. Trajectories were cued by displaying barriers and targets on the screen at various layouts, yielding initial movement directions and target directions that diverged by up to 90 degrees, in 45 degree increments (Fig. 3A). This resulted in 40 movement conditions (8 targets x 5 possible trajectories towards each target), which effectively disentangled initial and target directions. As before, this task followed an instructed-delay paradigm where a random subset of trials skipped the execution period (Fig. 3B).

**Fig. 3:**
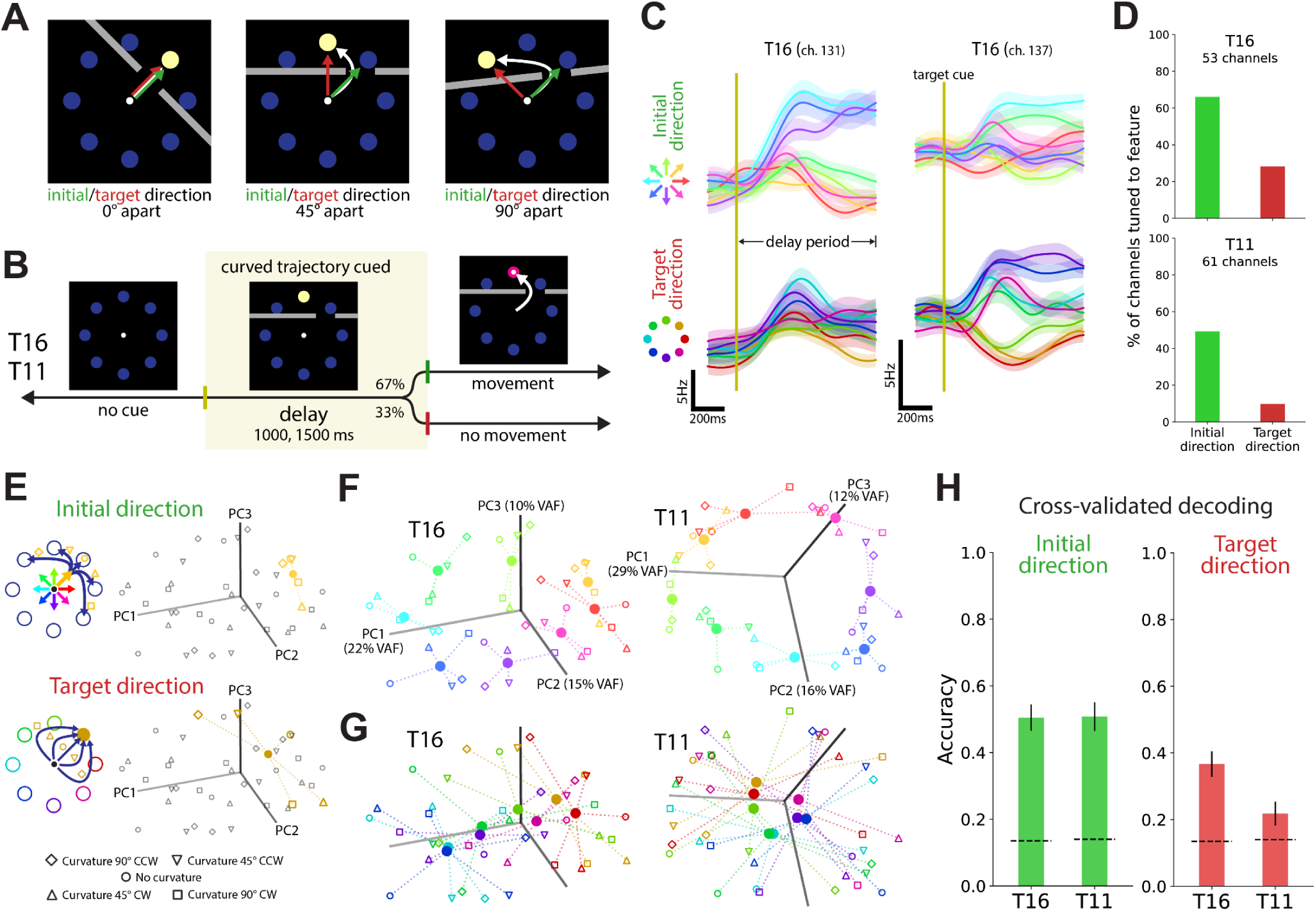
Preparatory activity is tuned to both the initial and endpoint directions of curved trajectories. **A.** Example on-screen layouts to cue movements with different directions and curvatures. **B.** Trial structure and timing for the instructed-delay task that cued the preparation and attempt of curved movements. **C.** Firing rates through time for example neural channels with trial-averages computed either across initial movement directions (top row) or target directions (bottom row). Shaded regions indicate +/-the standard error of the mean for each condition group. **D.** Percentage of active channels from each participant tuned to either the initial movement direction or target direction (*p*-value < 0.01, 2-way ANOVA main effects, Benjamini-Hochberg corrected). **E.** Neural population activity during the delay period (averaged across trials and time) for the different movement conditions (8 directions x 5 curvatures) as visualized by principal component analysis (top 3 components); markers indicate the corresponding curvature value for each neural activity projection. The top panel highlights neural projections for an example initial direction value and all corresponding movement curvature conditions, and the bottom panel is the same but for an example target direction instead. Dotted lines connect movements to different directions with the same curvature. Solid circle markers correspond to the average neural activity across curvatures for each direction. **F.** Same as E for both participants, but with all movement conditions colored according to initial movement direction. **G.** Same as E for both participants, but with all movement conditions colored according to target direction. **H.** Accuracy of single-trial predictions of initial movement direction and target direction decoded from preparatory activity during the delay period, using SVM classifiers (leave-one-out cross-validation). Error bars indicate the 95% confidence interval and dashed lines indicate chance levels.

At the level of individual neural channels, preparatory activity showed distinct tuning to the initial and target directions. For certain channels, neural activity was only discernible across movements with different initial directions, whereas other channels showed stronger modulation to the target direction (Fig. 3C). Aggregating across the population, 49-66% of active channels from each participant had preparatory activity tuned to the initial movement direction, compared to 10-28% being tuned to the target direction (*p*-value < 0.01, 2-way ANOVA main effects, Benjamini-Hochberg corrected; Fig. 3D). This indicates preparatory activity encodes both of these features from the upcoming movement trajectory, with stronger encoding of the initial direction compared to the target direction.

To visualize the relative strength of the preparatory activity tuning to the initial and target directions at the population level, we used principal component analysis (PCA) to compute the top dimensions of the neural population activity that explained the highest amount of variance during the delay period across all conditions. We then projected the neural population activity from each of the 40 different movement conditions onto these dimensions (Fig 3E). For both participants, the organization of activity on these top PCA dimensions was consistent with the initial direction of movement, even for movements with different curvatures (Fig. 3F). In contrast, the preparatory activity in these representative dimensions was not as structured across movements with the same target direction but with different curvatures (Fig. 3G). This again suggests that for curved trajectories, the movement feature that is more prominently encoded in preparatory activity is the initial movement direction.

To evaluate the applicability of these features for BCI applications, we tested how accurately the initial and target directions of upcoming movements could be predicted from preparatory activity on individual trials. Using SVM classifiers, we found that both features could be decoded with accuracy well above chance (chance: 14%; Fig. 3H), and with significantly higher accuracies for initial directions than for target directions (51% > 37% for T16, 51% > 22% for T11; *p*-value = 4.4 x 10^-7^ and 2.1 x 10^-22^ respectively, two-proportion z-test). These analyses confirm that preparatory activity encodes multiple features of curved movement trajectories, with stronger relative tuning to the initial direction compared to the trajectory target position.

### Preparatory activity is slightly tuned to upcoming movement distance and speed

An additional parameter of upcoming movement that is critical for practical brain-computer interface applications is the extent or distance of movement. If both direction and distance are encoded by preparatory activity, it could enable neural decoders that predict a movement’s endpoint before the movement is actually initiated. However, when probing tuning to movement distance, it is important to consider that distance and speed are typically correlated: movements to further distances are usually performed at higher speeds^38,39^. In previous studies of preparatory activity with non-human primates performing reaches, speed and distance have been challenging to tease apart^22^, though there is evidence of some degree of independence in their tuning^20^.

To separately evaluate preparatory tuning to distance and speed in human motor cortex, we developed a new instructed-delay task (inspired by previous work^40^) that explicitly cues the spatiotemporal properties of movement trajectories rather than spatial properties alone. In each trial, participants attempted to follow side-to-side movements of a “Pac-Man” icon, with the movement trajectory shown in advance as a dotted path indicating where Pac-Man would be at any point in time (Fig. 4A). Position and time were represented by the horizontal and vertical axes, respectively. The cued trajectories could have independently defined directions (dots spread to the right or left), distances (lateral extent), or speeds (slope), which allowed these movement parameters to be decoupled across trials. Consistent with an instructed-delay paradigm, dots were stationary during a delay period to allow preparation, and then scrolled downwards at a constant rate during an execution period (Fig. 4B). Movements were randomly selected from 18 total conditions, resulting from the possible combinations of two directions (right or left), three distances (short, medium, long), and three speeds (slow, medium, fast).

**Fig. 4:**
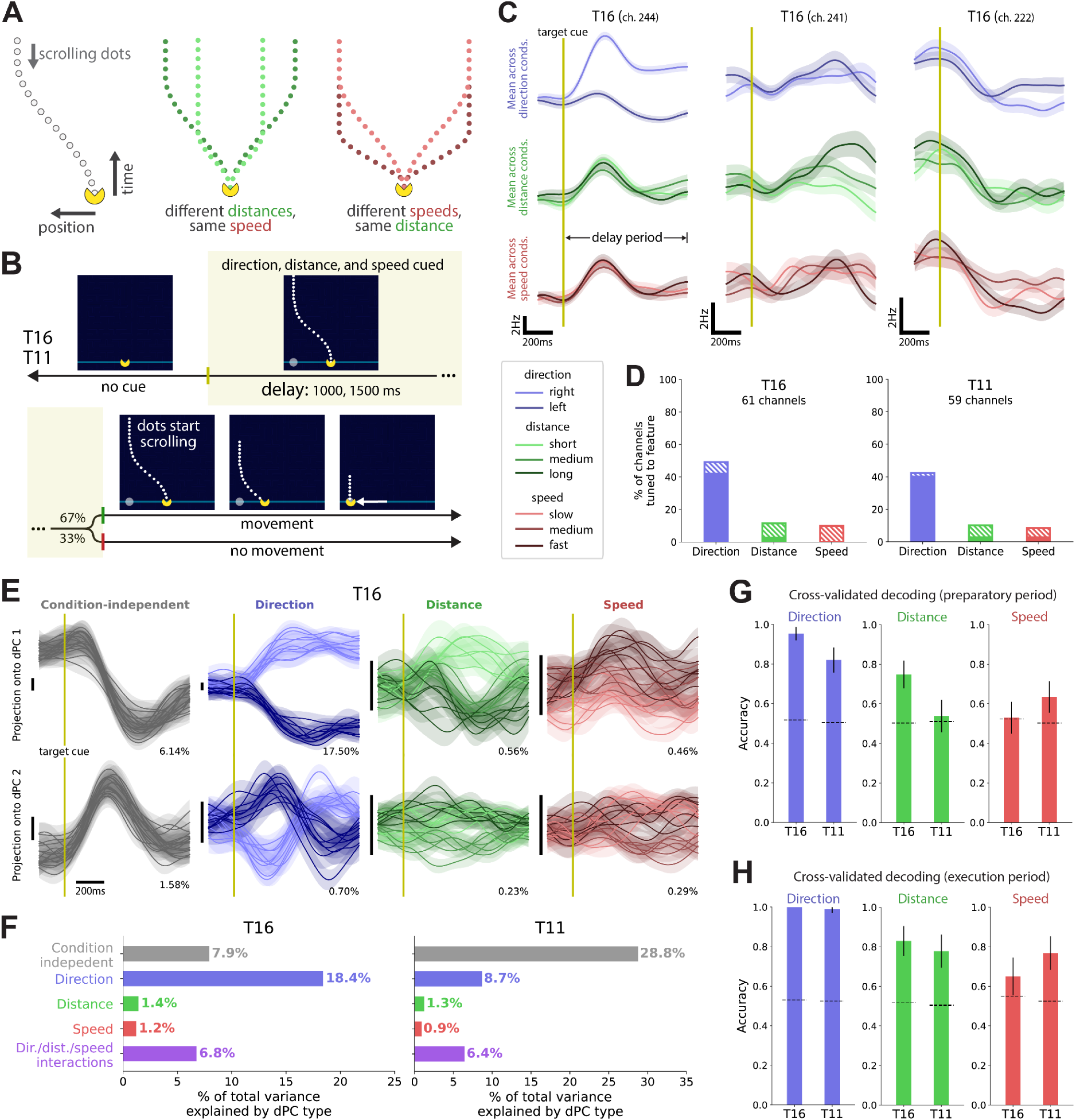
Preparatory activity is slightly tuned to upcoming movement distance and speed. **A.** Example scrolling dot trajectories used in the Pac-Man instructed-delay task to independently cue movement direction, distance, and speed. **B.** Trial structure and task timing: dots were stationary during the delay period, and scrolled at a constant speed during the execution period. **C.** Firing rates through time for example neural channels with trial-averages computed either across movement directions (top row), distances (middle row), or speeds (bottom row). Shaded regions indicate +/-the standard error of the mean for each condition group. **D.** Percentage of active channels from each participant tuned to either the direction, distance, or speed of movement (*p*-value < 0.05, 3-way ANOVA, Benjamini-Hochberg corrected). Solid bar indicates percentage of channels with a main effect of the specified movement feature and the hatched bar indicates percentage of channels with either a main effect or interaction with other features. **E.** Projection of average neural population activity during the delay period (for participant T16) onto demixed principal components (dPCs) that are condition-independent or specific to upcoming movement direction, distance, or speed. Traces are colored according to the value of the movement feature tied to each specific dPC and shaded regions indicate +/-the standard error of the mean for projections corresponding to trials from each condition. Vertical black bars to the left of each subplot indicate a consistent scale across dPC projections (arbitrary units). The number on the bottom right of each subplot indicates the percentage of total preparatory activity variance explained by each component. **F.** Percentage of the total trial-averaged preparatory activity variance accounted for by each dPC type. **G.** Accuracy of single-trial predictions of movement direction, distance, and speed decoded from preparatory activity during the delay period, using SVM classifiers (leave-one-out cross-validation). Only trials from the short/long distance conditions and slow/fast speed conditions were considered for this analysis. Error bars indicate the 95% confidence interval and dashed lines indicate chance levels. **H.** Same as G, but decoding from a window of neural activity during the execution period.

We first visualized individual neural channels’ mean preparatory activity across movements with different directions, distances, or speeds and found that some channels are selectively tuned to each of these features (Fig. 4C). In line with previous results, a substantial percentage of active channels – ranging from 40-43% across participants – showed tuning to upcoming movement direction across different distances and speeds (*p*-value < 0.05, 3-way ANOVA, Benjamini-Hochberg corrected; Fig. 4D). The proportion of channels tuned to movement distance or speed was notably lower: <4% had a main tuning effect for distance and <4% for speed. The range was slightly higher when interactions with the other features were included: 10-12% of active channels for distance and 8-10% for speed.

We then tested the degree to which representations of distance, direction, and speed were linearly separable in the neural population preparatory activity by using dPCA to decompose activity into components specific to each of these variables and their interactions (Fig. 4E, Supplemental Fig. 4). Consistent with the single-channel findings, the main components of preparatory activity (in terms of variance accounted for) were either tied to movement direction or condition-independent. The dPCs for distance or speed also showed some modulation during the delay period, but substantially smaller than for direction: distance and speed tuning each only accounted for ∼1% of the total preparatory activity variance in each participant, compared to the 9-18% variance accounted for by direction tuning (Fig. 4F). Additionally, 6-7% of the population activity variance was explained by the interaction of direction, distance, and speed, suggesting an integrated encoding of these task features in preparatory activity.

To verify that preparatory encoding of distance and speed is robust for individual trials, we fit SVM classifiers to separate preparatory activity between trials of short and long distances, slow and fast speeds, and both directions. As expected, movement direction could be reliably decoded from single-trial preparatory activity with accuracies ranging from 82-95% (chance: 52%; Fig. 4G). Decoding performance for distance and speed was more nuanced, with differences across participants: T16 had above-chance decoding of distance (accuracy = 75% > chance = 50%) but not speed, whereas the opposite was the case for T11 (accuracy = 63% > chance = 50%). This participant-specific low discernibility of speed or distance was not simply due to the movement attempts themselves being indistinguishable: when we repeated the same decoding analysis for neural activity during the execution period, decoding performance was well above chance for all conditions in both participants (Fig. 4H). Altogether, these decoding results confirm that even on single trials, human preparatory activity can be informative of upcoming movement distance and speed in addition to direction. The participant-specific reliability of this encoding may tie to differences in the precise placement of electrode arrays or in the strategy each participant uses to prepare for upcoming movement attempts.

### Online decoding from preparatory activity enables rapid and fluid brain-computer interface control

Given that preparatory activity in the human motor cortex is informative about features of upcoming movements, it should be possible for a BCI to decode a user’s intended movement trajectory using preparatory activity alone in online settings. Unlike standard BCI paradigms that rely on mapping neural activity during movement onto instantaneous control commands (Fig. 5A), a paradigm leveraging preparatory activity could precompute and execute trajectories quickly and fluidly as soon as the intention to initiate movement is detected. Along these lines, previous work with non-human primates has shown that predicting movement goals from preparatory activity can enable fast and accurate BCI control performance^30–32^.

**Fig. 5:**
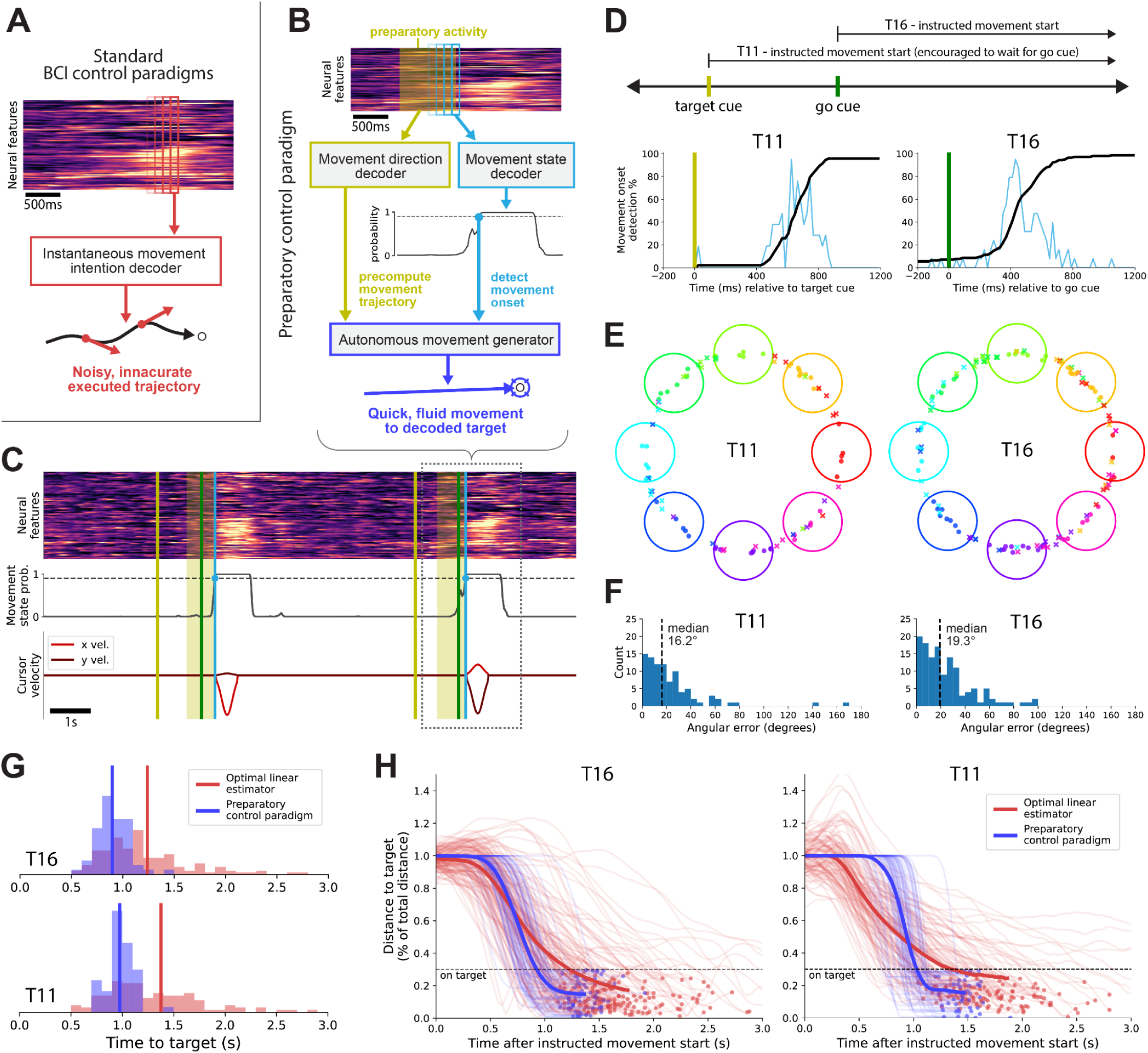
Online decoding from preparatory activity enables rapid and fluid brain-computer interface control. **A.** Standard BCI control paradigms rely on decoders that map execution-related neural activity onto instantaneous predictions of intended movement commands, which often produce inaccurate movement trajectories due to the characteristic moment-to-moment variability in neural activity. **B.** Schematic of self-paced online BCI control paradigm that decodes intended movements from preparatory activity. **C.** Example time window with normalized neural features, decoded movement state probability, and cursor velocity generated by the preparatory control paradigm, for participant T16. Yellow lines, green lines, and light blue lines indicate target cue times, go cue times, and decoded movement onset times, respectively. Yellow boxes indicate the windows of preparatory activity used to decode intended movement direction. **D.** Distribution of movement onset times decoded during online evaluation, relative to the target cue time for participant T11 or go cue time for participant T16. Light blue lines correspond to the histogram of decoded movement onset times and black lines are cumulative distributions. **E.** Predicted movement directions during online evaluation using the preparatory control paradigm. Large circles illustrate target positions cued to the participants during a radial-8 evaluation task. Small markers show predicted movement directions for each trial (with radial jitter added for clarity), colored according to the corresponding cued target directions. Dots indicate successful trials whose decoded movement ended at a distance to the target of up to 30% of the total distance from the start position to the target, whereas crosses indicate trials where movement ended further away. **F.** Distributions of angular errors in decoded movements (relative to the center of cued targets) for all trials during online evaluation with each participant. **G.** Time to target for successful trials of the radial-8 task using either an optimal linear estimator (OLE) or the preparatory control paradigm for control. Vertical lines indicate the mean time to target for each control scheme. **H.** Distance to target (as a percentage of the total distance from the start position to the target) as a function of time for successful trials using an OLE or the preparatory control paradigm for control. Each thin trace corresponds to a single trial and thicker traces correspond to the average across control schemes. Dots indicate the target acquisition time, which is computed for each trial as the time when the cursor has been on target for 500ms.

As a proof-of-concept of the use of preparatory activity in a self-paced BCI, we designed a preparatory cursor control paradigm with three components (Fig. 5B). The first component, a movement initiation decoder, continuously reads incoming neural activity features and outputs the probability of being in a “movement” state or not. We take movement initiation as the timepoint when the decoded probability crosses a predetermined threshold. This triggers the second component, a movement direction decoder, which predicts the intended movement direction from a window of preparatory activity preceding the detected movement initiation event. The last component, an autonomous movement generator, executes a quick cursor movement to the decoded movement goal position (at a fixed distance in the decoded direction). Of note is that the decoded direction can take any continuous value, making our control paradigm more versatile than previous non-human primate preparatory BCIs that only allowed movements to a discrete number of positions^30–32^.

To evaluate online performance, participants performed a modified version of the instructed-delay radial-8 task in which cursor movements were controlled by the preparatory control paradigm. As before, one of eight targets was presented in each trial, and following a brief delay, a go cue was shown to instruct a quick movement attempt toward the target. Notably, movement initiation could be decoded before or after the go cue and executed movements could end outside of the targets since direction decoding was not constrained to the 8 target positions. For T16, a percentage of trials omitted the go cue, and trials would fail if movements were initiated outside of the cued execution period. For T11, a timer indicating the remaining delay time was shown and he was instructed to attempt to match the timing of the go cue, though initiating movement early would not fail trials.

Participants successfully used the preparatory control paradigm to perform the task (Fig. 5C), achieving responsive detection of movement initiation and quick cursor movements that resembled able-bodied movements (400-600 ms)^38^. To evaluate the preparatory control paradigm’s ability to accurately infer intended movement timing, we recorded residual forearm EMG activity from participant T16 as a proxy for intended movement. Movement initiation inferred from neural and EMG activity were highly consistent (Supplemental Fig. 5), confirming the responsiveness of the movement state decoder. Responsiveness was also evident from the distributions of decoded movement onset times for both participants: ∼400 ms after the go cue for T16 and ∼600 ms after the target cue for T11 (Fig. 5D), both reasonable values for reaction time (plus some additional preparation time for T11 in attempting to match the timing of the go cue). In terms of directional accuracy, decoded directions mostly clustered around the corresponding cued target (Fig. 5E). The success rate, which we define as the percentage of trials where movement ended within a target radius of 30% of the total distance from start to target, was 45% for T16 and 56% for T11, which is promising for the simple implementation tested here without any capabilities to correct for decoding errors following movement initiation. Angular errors between the predicted movement directions and directions of center of the cued targets were also generally low, with median values of 19° for participant T16 and 16° for participant T11 (Fig. 5F).

Finally, we compared the performance of the preparatory control paradigm to that of an optimal linear estimator (OLE), a commonly used BCI control paradigm that continuously predicts intended movement velocity^41–43^. For successful trials, decoded movements reached the target much faster for the preparatory control paradigm than the OLE (mean time to target of 0.90 < 1.23 s for T16, 0.97 < 1.38 s for T11; *p*-value = 1.3 x 10^-8^ and 3.3 x 10^-7^ respectively, Mann-Whitney *U* test; Fig. 5G). Additionally, the time course of the distance between the cursor and target was markedly more consistent for the preparatory control paradigm (Fig. 5H). Trajectories could also be considered more efficient for this paradigm: the distance to target monotonically decreased as movement progressed, compared to the variability with the OLE control which at times increased the distance between the cursor and target. It is also worth noting that even with the enforced delay prior to movement which was the case during evaluation of the preparatory control paradigm with participant T11, it still allowed acquiring targets faster than the OLE. Overall, this proof-of-concept control paradigm implementation highlights the performance benefits of leveraging preparatory activity for BCI control applications.

## Discussion

Preparatory activity in the human motor cortex serves as a rich and underexplored source of information for neurally controlled assistive devices. Our work with clinical trial participants with tetraplegia demonstrates that preparatory activity is highly informative about many features of upcoming movements, including movement direction, effector, trajectory curvature, and to some extent, distance and speed. Furthermore, preparatory activity is robust enough to enable online BCI control, here allowing substantially faster cursor control than standard BCI algorithms. Altogether, our results highlight the potential of leveraging the rich information content in preparatory activity for practical clinical BCI applications.

### Preparatory activity reflects multiple levels of movement abstraction

In non-human primates, there is evidence for preparatory activity encoding abstract task-related movement features, such as the positions of targets regardless of physical movement direction^44^. Our results further demonstrate that abstract task-related features are also encoded by preparatory activity in the human motor cortex, through preserved directional tuning across movements with different effectors as well as preserved endpoint direction tuning for movements following different curved trajectories. Preparatory activity was also comparable between tasks with different saliency in visual stimuli, indicating this abstract task-related tuning was not purely due to encoding of visual task features. Separately, a substantial part of preparatory activity was linked to features closely related to motor execution, such as the effector identity and initial movement direction. Together, these findings indicate preparatory activity encodes a mixture of abstract and movement-specific variables, with the potential role to anticipate neural computations that transform abstract features from upstream brain areas into muscle-like motor commands^37^.

### Strength of preparatory activity tuning to movement distance and speed compared to direction

Our results show that preparatory activity in human participants encodes the distance and speed of movement, but this encoding only accounts for a fraction of the total preparatory modulation. This is at odds with what is expected from studies in non-human primates, where preparatory tuning to movement distance and/or speed is more prevalent and accounts for a substantial part of the population activity^20,22^. In this regard, an important consideration with clinical trial participants is they do not receive proprioceptive or visual feedback during task performance due to the lack of overtness in attempted movements. Without feedback, more variability is expected during movement execution and its associated preparation. Movements with different distances and speeds are also tied to a grading in the intensity of attempted muscle activations, which might be perceived similarly during preparation. Comparatively, movements towards different directions involve distinct sets of muscles, with motor plans that are easier to distinguish.

### Rich preparatory encoding as a mechanism for optimal generation of movement

The encoding of diverse movement features we observed is likely a limited view of a much more complex and less interpretable neural state that arises during motor preparation. In the framework of neural computations through population dynamics^45^, preparatory activity can be seen as an optimal initial condition for the generation of upcoming movement^14,15^. Our findings are consistent with this framework, since the distinct preparatory activity we observed for diverse movement features can be interpreted as the corresponding initial neural state for each movement condition. From this perspective, each movement that is perceived and prepared differently would have its unique preparatory neural state. Further investigation would require scaling task complexity to explore an even greater diversity of movements, and ideally, an increase in recording channels to appropriately account for the information encompassed by the increasing number of movement conditions.

### Implications for brain-computer interface control applications

In this work we also showed, for the first time with clinical trial participants, that anticipated predictions of intended movement decoded from preparatory activity can effectively drive BCI control of a computer cursor. Even with an overly simplistic implementation, the proof-of-concept control scheme we tested enabled substantially faster movements than standard methods. Decoded movements were also not affected by moment-by-moment neural noise, which could prove to be less cognitively demanding for users. Further exploration of this type of control scheme could be a crucial step towards closing the gap between current BCI control performance and that of able-bodied movement.

Our characterization of preparatory activity across different tasks also highlights potential future challenges for preparatory control paradigms. The variability in single-trial tuning can lead to inaccurate predictions of intended movement trajectories, which could require online detection and correction of motor error signals^46,47^. Encoding of movement distance might also not prove robust enough for pre-programming trajectories that can reach any point in space. As an alternative, a paradigm which generates fast precomputed movements in a specific direction but detects the transient intention to stop movement at a specific distance^48,49^ – instead of decoding the intended distance in advance – might prove more practically feasible.

## Acknowledgments

The authors thank participants T16, T11, T5, and their families and care partners for their contributions to this research. The authors also thank Yvan Bamps, Haris Rashid, Colleen Spellen, Maryam Masood, and Dave Rosler for administrative support and clinical research support.

This work was supported by NIH-NINDS/OD DP2NS127291, Simons Foundation as part of the Simons-Emory International Consortium on Motor Control (CP), NIH F32HD112173 (SRN), NIH-NIDCD U01DC017844, Department of Veterans Affairs Rehabilitation Research, Development, and Translation A2295R, A4820R (LRH), NIH-NIDCD R01DC014034, Simons Foundation Collaboration on the Global Brain 543045, Larry and Pamela Garlick, Wu Tsai Neurosciences Institute (JMH), and Howard Hughes Medical Institute (DTA). The content is solely the responsibility of the authors and does not necessarily represent the official views of the National Institutes of Health, or the Department of Veterans Affairs, or the United States Government.

## Author contributions

MR and CP conceptualized the project. MR designed the tasks and research sessions, implemented the online preparatory control paradigm, and performed data analyses. MR, PB, BR, and SRN conducted research sessions with participant T16. MR, AA, and CN conducted research sessions with participant T11. MR and NH conducted research sessions with participant T5. DA and MR deployed the hardware and software required to run research sessions with participant T5. SRN and YA led research sessions where data for online cursor control using an optimal linear estimator was collected with participants T16 and T11. SRN, MR, and YA contributed to software development to support research sessions. LRH is the sponsor-investigator of the multi-site clinical trial. LRH supervised research with participant T11 and JMH supervised research with participant T5. CP and NAY supervised research with participant T16 as well as all aspects of the project. Funding was acquired by CP, SRN, LRH, and JMH. MR and CP wrote the manuscript. All authors reviewed and contributed to edits to the manuscript.

## Declaration of generative AI and AI-assisted technologies in the writing process

During the preparation of this work the authors used ChatGPT (GPT-4o/GPT-5) in order to improve readability and language of the manuscript. After using this tool, the authors reviewed and edited the content as needed and take full responsibility for the content of the published article.

## Methods

### Study participants

This study includes data from three participants, referred to as T16, T11, and T5, who each gave informed consent and were enrolled in the BrainGate2 Neural Interface System clinical trial (ClinicalTrials.gov Identifier: NCT00912041, registered June 3, 2009). This pilot clinical trial was approved under an Investigational Device Exemption (IDE) by the US Food and Drug Administration (Investigational Device Exemption #G090003; CAUTION: Investigational device. Limited by federal law to investigational use), as well as granted approval by the Emory University IRB (protocol #00003070), the Mass General Brigham IRB (protocol #2009P000505), the Providence VA Medical Center, and the Stanford University IRB (protocol #20804). All research was performed in accordance with relevant guidelines and protocols.

T16 is a right-handed woman, 52 years of age at the time of data collection for this study, with tetraplegia and dysarthria due to a pontine stroke which occurred approximately 19 years prior to enrollment in the BrainGate2 clinical trial. T16 had four 64-channel intracortical microelectrode arrays (1.5 mm electrode length; Blackrock Neurotech, Salt Lake City, Utah, USA) placed in her left precentral gyrus, guided by individualized Human Connectome Project (HCP) cortical parcellation: two in HCP-identified hand knob area (6d), one in HCP-identified ventral premotor cortex (6v), and one on the border of the HCP-identified premotor eye fields (PEF) and speech-related area 55b. T16 is able to speak slowly and quietly, but enunciation is restricted by limited face and mouth movement. She has limited voluntary control of her upper extremities, with some shoulder motion and some slow and contractured wrist and finger movements. She has limited to no voluntary control of her lower extremities. T16’s sensation is fully intact. Data for this study was collected with T16 between post-implant days 253-543.

T11 is a right-handed man, 38 years of age at the time of data collection for this study, with tetraplegia due to a cervical spinal cord injury (classified as C4 AIS-B) which occurred approximately 11 years prior to enrollment in the BrainGate2 clinical trial. T11 had two 96-channel intracortical microelectrode arrays (1.5 mm electrode length; Blackrock Neurotech) placed in his left dorsal precentral gyrus, targeting the hand knob area (6d) as identified anatomically by preoperative magnetic resonance imaging (MRI). T11 has full movement of his face and head with limited voluntary motion of the arms. Data for this study was collected with T11 between post-implant days 1321-1961.

T5 was a right-handed man, 70 years of age at the time of data collection for this study, with tetraplegia due to cervical spinal cord injury (classified as C4 AIS-C) which occurred approximately 9 years prior to enrollment in the BrainGate2 clinical trial. T5 had two 96-channel intracortical microelectrode arrays (1.5 mm electrode length; Blackrock Neurotech) placed in his left dorsal precentral gyrus, targeting the hand knob area (6d) as identified anatomically by preoperative MRI. T5 had full movement of his face and head and the ability to shrug his shoulders. Below the level of spinal cord injury, T5 had limited voluntary motion of the legs and arms. Data for this study was collected with T5 during post-implant day 2623.

### Research sessions and data collection

For each participant, data was collected during research “sessions” that took place across different days. All research sessions were performed at the participant’s place of residence. During each session, the participant was positioned at an idle position in front of a computer monitor (screen size: between 24 and 24.5 inches diagonal, 1920 x 1080 pixels), and asked to perform different motor tasks in a series of “blocks” lasting 3-6 minutes each. Table 1 lists all research sessions and number of blocks during which data was collected with each participant.

**Table 1:** Research sessions with participants.

| Participant | Session (year-month-day) | Data collected |
| --- | --- | --- |
| T16 | 2024-08-16 | Open-loop instructed-delay radial-8 task using attempted wrist movements (6 x 4 min blocks, 382 trials; Fig. 1) and attempted finger movements (6 x 4 min blocks, 381 trials; Fig. 2), open-loop instructed-delay Pac-Man task using attempted wrist movements (6 x 4 min blocks, 345 trials; Fig. 4) |
|  | 2024-10-02 | Closed-loop instructed-delay radial-8 task using OLE (2 x 5 min blocks, 81 trials; Fig. 5) |
|  | 2024-10-23 | Closed-loop instructed-delay radial-8 task using OLE (2 x 5 min blocks, 81 trials; Fig. 5) |
|  | 2024-11-11 | Closed-loop instructed-delay radial-8 task using OLE (3 x 5 min blocks, 133 trials; Fig. 5) |
|  | 2025-02-28 | Open-loop instructed-delay curved radial-8 task using attempted wrist movements (11 x 4 min blocks, 622 trials; Fig. 3) |
|  | 2025-04-25 | Open-loop instructed-delay radial-4 task using attempted wrist movements with regular visuals (3 x 4 min blocks, 190 trials) and suppressed visuals (3 x 4 min blocks, 186 trials; Supplemental Fig. 2) |
|  | 2025-06-02 | Closed-loop instructed-delay radial-8 task using preparatory control paradigm (3 x 4 min blocks, 198 trials; Fig. 5) |
| T11 | 2023-05-07 | Closed-loop radial-8 task using OLE (2 x 4 min blocks, 227 trials; Fig. 5) |
|  | 2023-06-18 | Closed-loop instructed-cued-delay radial-8 task using preparatory control paradigm (1 x 4 min block, 93 trials; Fig. 5) |
|  | 2025-01-13 | Open-loop instructed-delay radial-8 task using attempted wrist movements (4 x 4 min blocks, 264 trials; Fig. 1) and attempted finger movements (6 x 4 min blocks, 264 trials; Fig. 2), open-loop instructed-delay Pac-Man task using attempted wrist movements (6 x 4 min blocks, 326 trials; Fig. 4) |
|  | 2025-02-05 | Open-loop instructed-delay curved radial-8 task using attempted wrist movements (9 x 4 min blocks, 514 trials; Fig. 3) |
| T5 | 2023-10-23 | Open-loop instructed-cued-delay radial-8 using attempted wrist movements (3 x 4 min blocks, 335 trials; Fig. 1) |

Data collection was run on a dedicated computer rig. Execution and communication of modular Python and C++ processes for running tasks and synching with neural recordings were handled using the Backend for Realtime Asynchronous Neural Decoding system (BRAND)^50^.

### Intracortical neural signal acquisition and processing

Intracortical electrical voltages were recorded from microelectrode arrays using NeuroPort Neural Signal Processor systems (Blackrock Neurotech). Recorded signals were first analog band-pass filtered (0.3 Hz to 7.5 kHz, 4th order Butterworth filter) and digitized at 30 kHz (250 nV resolution). Linear regression referencing (LRR)^51^ was then applied, with electrode-specific LRR coefficients determined for each block using the filtered 30 kHz data either from the same block for offline analyses (Figs. 1-4, Supplemental Figs. 1, 3, 4), or from data collected during the previous block for online decoding results (Fig. 5, Supplemental Fig. 5). For offline analyses, the resulting voltage time series were then digitally band-pass filtered (250 Hz to 5000 Hz, acausal 4th order Butterworth filter). For online decoding results, voltage time series were digitally high-pass filtered (>250 Hz, acausal 4th order Butterworth filter with 4 ms delay).

After rereferencing and filtering, threshold crossings were computed for each neural channel. Thresholds were set to-4.5 times the root-mean-square of the voltage signal for each channel (except for the 2023-05-07 and 2023-06-18 sessions with participant T11, where the multiplier was-3.5), computed on the same block for offline analyses or from a previous block for online decoding. Threshold crossings were then grouped into bins with a duration of 10 ms (2023-05-07 and 2023-06-18 sessions with participant T11, Fig. 5) or 20 ms (all other results). Spike-band power (SBP) was also computed for each channel, by taking the mean of squared values of the rereferenced and filtered 30 kHz samples in each time bin. Only channels from arrays in the hand-knob area of participants’ precentral gyrus (128 channels for T16, 192 channels for T11 and T5) with a mean firing rate of at least 2 Hz across all time points and blocks for each particular task (denoted “active” channels) were considered for analysis or online decoding. Binned threshold crossings and spike-band power for active channels were then appended to form a vector of “neural features” for each time bin. Finally, neural features were smoothed acausally using a Gaussian kernel with σ = 100 ms for offline analyses or causally using exponential smoothing for online decoding (smoothing parameter varied across sessions, see below).

For neural population analyses (all analyses except peri-stimulus time histograms and single-channel tuning), binned neural features were z-scored (each channel’s activity was baseline-centered and divided by its standard deviation) to prevent channels with high magnitude values from dominating the variance of the data. For offline analyses, the baseline subtracted from each channel’s values was computed on a per-block basis (to account for baseline shifts across the session) and the standard deviation was computed across all blocks for a particular task (after baseline-centering). For each block of online decoding, the baseline and standard deviation for z-scoring was computed on the previous block.

### Electromyography activity acquisition

Electromyography activity for participant T16 was recorded using a Trigno Duo sensor (Delsys Inc., Natick, Massachusetts, USA). Electrical voltages were recorded from electrodes placed on the ventral and dorsal surfaces of T16’s forearm (2 channels total). Recorded signals were first analog band-pass filtered (100 Hz to 850 Hz, Butterworth filter with 2-pole high-pass corner and 4-pole low-pass corner) and digitized at 2148 Hz (168 nV resolution). The resulting voltage time series were then digitally band-pass filtered (100 Hz to 500 Hz, acausal 4th order Butterworth filter), squared to compute the signal power, and digitally low-pass filtered (<5 Hz, acausal 4th order Butterworth filter). To align with the neural spiking data, the processed EMG activity samples were subsequently resampled to 100 Hz (through linear interpolation) and grouped into 20 ms bins (by averaging across samples).

For plotting EMG activity (Supplemental Figs. 2, 5), each channel’s samples were baseline-centered and divided by their standard deviation. The baseline subtracted from each channel’s values was computed on a per-block basis and the standard deviation was computed across all blocks for a particular task (after baseline-centering).

### Eye tracking data collection

Gaze position data for participant T16 was collected using a Tobii Pro Spark eye tracker (Tobii AB, Stockholm, Sweden). Data was collected at a sampling rate of 60 Hz. To align with the neural spiking data, eye tracking samples were resampled to 100 Hz (through linear interpolation) and grouped into 20 ms bins (by averaging across samples).

### Instructed-delay tasks

During research sessions, participants performed tasks with an instructed-delay paradigm, structured as a sequence of “trials”. Each trial began with a “target cue” marking the start of a delay period, during which participants were instructed not to move while being visually cued on the target position and features of the upcoming movement. Some tasks also showed a circular timer during the delay period to indicate the remaining delay time (‘instructed-cued-delay’). The delay duration and cued movement condition were selected through pseudorandom sampling from a distribution promoting a balanced number of trials for each combination of these values within each block. The delay period was followed by an execution period indicated by a “go cue” where participants attempted to match quick on-screen cursor movements consistent with what was previously cued and then remained on the target position for a specified hold time. After the execution period, the go cue turned off to indicate an inter-trial period where participants returned to a rest state. For some tasks, a random subset of trials skipped the go cue (33% probability), such that the delay period was immediately followed by the inter-trial period preceding the next trial. Participants performed different instructed-delay tasks designed to cue different movement features (Table 1): “radial-8” and “radial-4” tasks to cue direction, “curved radial-8” task to cue separate initial and target directions, and “Pac-Man” task to cue direction, distance, and speed. Tasks could have the cursor move automatically during the execution period to perform the cued movement – in which case participants were instructed to track the cursor without visual feedback of their own attempted movement (“open-loop”) – or have cursor movements be controlled online from recorded neural activity (“closed-loop”). Possible values for the delay period duration, hold times, and inter-trial times for each task are indicated in Table 2.

**Table 2:** Timing parameters for different tasks and participants.

| Task | Participant | Delay period duration | Target hold time | Inter-trial time |
| --- | --- | --- | --- | --- |
| Open-loop radial-8 | T16 | 1000, 1500 ms | 600 ms | 1500 ms |
|  | T11 | 1000, 1500 ms | 600 ms | 1500 ms |
|  | T5 | 200, 600, 1000, 1400 ms | 100 ms | 600 ms |
| Open-loop curved radial-8 | T16 | 1000, 1500 ms | 600 ms | 1500 ms |
|  | T11 | 1000, 1500 ms | 600 ms | 1500 ms |
| Open-loop Pac-Man | T16 | 1000, 1500 ms | 500 ms | 1500 ms |
|  | T11 | 1000, 1500 ms | 500 ms | 1500 ms |
| Closed-loop radial-8 (using preparatory control paradigm) | T16 | 800, 1200 ms | 800 ms | 2000 ms |
|  | T11 | 800, 1000 ms | 500 ms | 600 ms |
| Closed-loop radial-8 (using optimal linear estimator) | T16 | 4000 ms | 1000 ms | 0 ms |
|  | T11 | n/a | 500 ms | 0 ms |

For the open-loop instructed-delay radial-8 and radial-4 tasks, participants attempted movements to match on-screen cursor movements towards radially distributed targets. 8 or 4 targets were shown at a fixed distance (400 pixels) from the center of the screen with equal angular spacing. All targets were initially dark during the inter-trial period with the cursor centered on the screen. For each trial, a target was selected during the delay period and was highlighted in yellow. The target remained highlighted for the duration of the delay period for participants T16 and T11; for T5 the target turned dark again 200 ms into the delay period. During the execution period, the highlighted target turned red (or green in the case of T5) to signal the go cue. The cursor then moved automatically towards the target following a truncated Gaussian velocity profile, with movement durations of 1.1 s for T16, 0.8 s for T11, or 0.5 s for T5. Participants were instructed to match cursor movements using either their wrists (Fig. 1, 2, Supplemental Figs. 1-3) or fingers (small finger for participant T16 or index finger for participant T11; T5 did not perform the task using his finger; Fig. 2). Once the cursor reached the target, the target turned pink (or red in the case of T5). After this the trial ended and the cursor returned to the center, followed by an inter-trial period that preceded the next trial.

An alternate version of the open-loop instructed-delay radial-4 task with suppressed visuals was also run with participant T16 (Supplemental Fig. 3). For this version of the task, movement direction was cued during the delay period by an arrow within a yellow circle at the center of the screen, and this circle turned red to indicate the go cue. No cursor was shown on screen during this task. T16 was also given the additional instruction to try to maintain her gaze fixated on the center of the screen as best as possible while performing this task, and an additional center target was shown during the inter-trial period to provide a gaze fixation point.

For the open-loop instructed-delay curved radial-8 task, participants used attempted movements of their wrists to match on-screen cursor movements following curved trajectories (Fig. 3). As in the base radial-8 task, 8 targets were shown at a fixed distance (400 pixels) from the center of the screen with equal angular spacing. Targets were initially dark during the inter-trial period. For each trial, one of 8 possible targets and one of 5 possible trajectories to that target (where the initial trajectory direction diverged by up to 90 degrees in 45 degree increments) were selected. During the delay period, the target was highlighted in yellow and barriers were displayed halfway between the center of the screen and the target, with an opening in the direction of the initial trajectory direction. To signal the go cue, the highlighted target turned red and the cursor moved automatically towards the target following a curved trajectory that evaded the barriers, with a truncated Gaussian speed profile and duration that scaled with trajectory length (ranging from 1.2 s to 2.1 s for T16 or 0.8 s to 1.4 s for T11). The highlighted target turned pink when the cursor came in contact with it, after which the cursor returned to the center followed by an inter-trial period.

For the open-loop instructed-delay Pac-Man task, participants used attempted movements of their wrists to match side-to-side on-screen movements of a Pac-Man icon (Fig. 4). The Pac-Man began each trial centered with no targets shown. During the delay period, the direction (right or left), endpoint distance from the center (200, 400, or 600 pixels), and average speed (200, 400, or 600 pixels/s) of the upcoming movement were cued through a stationary trajectory of dots displayed on screen (which followed a truncated Gaussian velocity profile): position and time were represented by the horizontal and vertical axes, respectively. The target position was also highlighted by a gray vertical line and a gray circle in the horizontal Pac-Man axis. To signal the go cue, the dotted trajectory started scrolling downwards at a constant rate during the execution period and the Pac-Man automatically moved horizontally towards the target following the scrolling dots. After reaching the target and remaining in its position for a specified time, the trial ended and Pac-Man returned to the center position followed by an inter-trial period.

Participants also performed closed-loop variations of the radial-8 task where cursor movements were decoded online from neural activity either using a preparatory control paradigm or an optimal linear estimator (OLE) (Fig. 5, Supplemental Fig. 5). For the preparatory control paradigm blocks with participant T16, the task followed the instructed-delay paradigm and trials could be failed if movement was initiated before the go cue or after a timeout of 3 s when the decoded movement ended off-target, in which case the task skipped to the inter-trial period. For the preparatory control paradigm blocks with participant T11, the task followed the instructed-cued-delay paradigm and early movements before the go cue were not failed. OLE blocks with participant T16 also followed the instructed-delay paradigm. OLE blocks with participant T11 did not have a delay period, so he was instructed to move the cursor towards the target immediately after it was cued. For both participants’ OLE blocks, the cursor position was not automatically recentered at the end of each trail, so trials alternated between outwards and inwards movements (only outwards trials were considered for analyses).

### Peri-stimulus time histograms

Peri-stimulus histograms (PSTHs) for each channel during the different tasks were computed by averaging smoothed threshold crossings for all trials from each condition, aligned either to their respective target cue for delay period activity (Figs. 1-4, Supplemental Fig. 3) or go cue for execution period activity (Fig. 1). PSTHs of the delay period activity only considered trials with a delay duration of at least 1000 ms, meaning that trials from participant T5 with delay durations of 200 ms and 600 ms were excluded from analysis.

### Single-channel tuning analyses

For each task, channels with preparatory activity tuned to different movement features were identified using ANOVA. Specifically, separate ANOVA tests were performed on the delay period activity from each channel during each task. For all ANOVA tests, the dependent variable was the average single-channel firing rate during the delay period: between 400 and 1000 ms after the target cue over all trials for T16 and T11 and between 400 and 800 ms after the target cue for trials with a delay duration of at least 1000 ms for T5. For the radial-8 task data, 1-way ANOVAs were performed to identify channels where preparatory activity was affected by the upcoming movement direction (*p*-values < 0.01). For the curved radial-8 task data, 2-way ANOVAs were performed to identify channels where preparatory activity was affected by either the initial or target directions of the upcoming curved movement (main effect *p*-values < 0.01). For the Pac-Man task data, 3-way ANOVAs were performed to identify channels where preparatory activity was affected by either the direction, distance, or speed of upcoming movement (separately reported main effect and interactions *p*-values < 0.05). For the 2-way and 3-way ANOVAs, *p*-values were corrected for multiple comparisons using the Benjamini–Hochberg procedure^52^.

### Demixed principal component analysis

Demixed principal component analysis (dPCA)^36^ was used to decompose preparatory population activity from the different tasks onto distinct components specific to each of the different movement features (or independent to all features). dPCA was performed using publicly available code: https://github.com/machenslab/dPCA. The built-in procedure to determine the best regularization term to prevent overfitting (10-fold cross-validation) was used for the data from each task. For the single-effector radial-8 task data (Fig. 1), the features/marginalizations of interest were time (condition-independent) and direction (condition-dependent). For the multi-effector radial-8 task data (Fig. 2), the features/marginalizations of interest were time (condition-independent), direction (effector-independent), effector identity (effector-dependent), and direction/effector interactions (also effector-dependent). For the Pac-Man task data (Fig. 4), the features/marginalizations of interest were time (condition-independent), direction, distance, speed, and direction/distance/speed interactions. For the data from each task and participant, dPCA projection matrices were fit on activity in the window from 0 ms to 1000 ms after the target cue, except for radial-8 data with T5, where dPCA was fit over the window from 0 ms to 800 ms after the target cue for trials with a delay duration of at least 1000 ms.

### Cross-validated movement feature decoding from preparatory activity

Single-trial movement feature decoding from preparatory activity during the open-loop tasks was computed using cross-validated support vector machine (SVM) classifiers with a linear kernel. The input to the SVM for each trial was a neural feature vector constructed by concatenating the neural feature values (threshold crossings and SBP) across all time bins in the window from 400 ms to 1000 ms after the target cue (30 bins). For participant T5, only trials with a delay duration of at least 1000 ms were used for this decoding analysis, and only neural features up to 800 ms after the target cue (20 bins) were concatenated to construct the input feature vector. Classifiers were trained through leave-one-out cross-validation: for each fold, all trials except one were included for training and performance was evaluated on the held-out trial. To choose the SVM regularization parameters for each fold, a nested 10-fold cross-validation was performed within the training data. Decoding performance was then computed as the mean accuracy on held-out trials across all folds and chance was computed as the mean frequency of the most frequent class label in the training set across all folds.

Decoding analyses were performed separately on data from the different tasks. All available trials were used for decoding analyses on data from the radial-8, radial-4, and curved radial-8 tasks (Figs. 1-3, Supplemental Fig. 3). For data from the Pac-Man task, only trials with values corresponding to either short or long distances and slow or fast speeds were considered. Therefore, trials with a distance of 400 pixels or speed of 400 pixels/s were excluded from the decoding analysis.

### Cross-validated pairwise neural correlations

Pairwise correlations between preparatory activity for movements with different directions and/or effectors (Fig. 2) were computed using a cross-validated estimator that reduces bias^37^. This estimator can result in values with magnitude greater than 1, which should be interpreted as the true correlation magnitude being close to 1. Preparatory activity vectors used to compute pairwise correlations were obtained by averaging the normalized neural features (threshold crossings and SBP) across time in the window from 400 ms to 1000 ms after the target cue. The average activity across directions within each effector condition was subtracted prior to time-averaging and computing correlations, to compare directional tuning within the same effector-agnostic reference frame.

### Visualization of population activity using principal component analysis

Preparatory population activity during the curved radial-8 task was visualized by using principal component analysis (PCA) to compute projections onto the top-3 dimensions of activity that explained the highest amount of variance during the delay period across all conditions (Fig. 3). For each of the 40 conditions (all valid combinations of initial directions and curved movement trajectories), neural features (threshold crossings and SBP) were trial-averaged and had the mean activity across conditions subtracted. The resulting trial-averaged neural features were then averaged across time for the window from 400 ms to 1000 ms after the target cue, and concatenated to create a matrix with rows equal to the number of neural features and columns equal to the number of conditions. PCA was then applied on this matrix to compute an orthogonal linear transformation that projects the high-dimensional neural features onto a 3-dimensional space.

### Preparatory control paradigm for online cursor control

The first component of the preparatory cursor control paradigm was a movement initiation decoder. This decoder continuously read a stream of incoming normalized neural features (threshold crossings and SBP for participant T16, only threshold crossings for participant T11) and exponentially smoothed them (time constant of 170 ms for T16, 144 ms for T11). A linear discriminant analysis (LDA) classifier then computed the probability that these neural features corresponded to either a “movement” or “no movement” state. During online control, if the probability output by this LDA classifier crossed a threshold of 0.9 for T16 or 0.98 for T11, the controller state was switched to the movement state, which triggered the other components of the control paradigm.

The second component of this control paradigm was a movement direction decoder. This decoder continuously buffered samples (40 x 20 ms bins for T16, 20 x 10 ms bins for T11) from a stream of incoming normalized neural features and exponentially smoothed them (same time constants as the movement initiation decoder for both participants). In the case of participant T16, a linear transformation (computed using PCA) was then applied to reduce the smoothed neural features to a lower-dimensional vector. Once a switch to the movement state was detected by the movement initiation decoder, a linear decoder predicted the relative position of the upcoming target (in cartesian coordinates) from the processed buffered preparatory activity preceding the detected movement initiation time. The direction of intended movement was computed as the angle towards these predicted coordinates.

The last component of this control paradigm was an autonomous movement generator. When movement was detected by the movement initiation decoder and an intended movement direction was decoded from preparatory activity, this movement generator executed a quick cursor movement to the decoded movement goal position (at a fixed distance in the decoded direction). This autonomous movement followed a truncated Gaussian velocity profile, with a duration of 600 ms for T16 or 400 ms for T11. After reaching the decoded goal position, the cursor held in place until the next trial.

To train the preparatory control paradigm presented in this work, we first collected calibration blocks with each participant during the evaluation sessions: 3 x 3 min open-loop instructed-cued-delay radial-8 blocks with participant T11 and 5 x 4 min open-loop instructed-delay radial-8 blocks with T16. To fit the LDA classifier for the movement initiation decoder with T16, training samples in the window from 400 ms to 0 ms before the target cue and 400 ms to 1000 ms after the target cue were labeled as “no movement” and samples in the window from 400 ms to 1000 ms after the go cue were labeled as “movement”. For trials that skipped the go cue, samples in the window from 400 ms to 1000 ms after the end of the delay period were also labeled as “no movement”. To fit this LDA classifier with participant T11, training samples in the window from 400 ms to 200 ms before the go cue were labeled as “no movement” and samples in the window from 200 ms to 400 ms after the go cue were labeled as “movement”. The movement direction decoder was fit using ridge regression: the inputs to the model were the time-flattened samples of the preparatory activity buffer preceding the decoded movement initiation for each calibration trial and the output were the relative cartesian coordinates of the target position. A cross-validated hyperparameter sweep was run on the training data to determine the regularization value for the ridge regression. For participant T16, the linear transformation to reduce neural data dimensionality prior to movement direction decoding was computed by using PCA to find the components of activity that explained ≥90% of the preparatory activity variance during the calibration blocks.

### Optimal linear estimator for online cursor control

Online cursor control using optimal linear estimators (OLEs) was used as a performance comparison to the preparatory control paradigm. The OLE decoders continuously predicted a participant’s intended cursor velocity from a linear readout of an incoming stream of normalized neural features (threshold crossings and SBP). For participant T16, neural features were exponentially smoothed (time constant of 90 ms) before being input to the decoder. For both participants, the velocity predictions from the decoder were also exponentially smoothed (time constant between 28 ms and 45 ms for T11, 70 ms for T16). The OLE implementation used with participant T16 included a softplus nonlinearity in its speed output (to suppress movement at low intended velocities). The OLE control data with participant T11 has been partially released in previous work^50^.

OLE decoders were trained on calibration blocks that preceded evaluation. For participant T16, calibration blocks consisted of 6 mins of the radial-8 task: 1 min of open-loop control where T16 attempted to match on-screen cursor movements to different targets followed by 5 mins of closed-loop control where cursor movements were driven by an OLE decoder. An initial decoder to map neural features onto intended cursor velocities was fit using ridge regression on data from the open-loop portion. New decoders were then fit every 5 s (similar to procedures for decoder calibration from previous studies^53,54^) using up to the previous 5 mins of open-loop and closed-loop data. For data from the closed-loop portion used for training, intended cursor velocities at every timepoint were assumed to be the unit vector from the cursor to the target scaled by the magnitude of the original decoded velocity, or zero if the cursor was on the target. To ease control during the start of the closed-loop portion of the task, components of decoded cursor velocities that were perpendicular to a straight line from the cursor to the target were attenuated, with attenuation decreasing from 100% to 0% over the course of the block. For participant T11, the calibration block consisted of 3 min of the open-loop radial-8 task, and the OLE decoder was fit using ridge regression on data from the whole block.

## Supplemental figures

**Supplemental Fig. 1:**
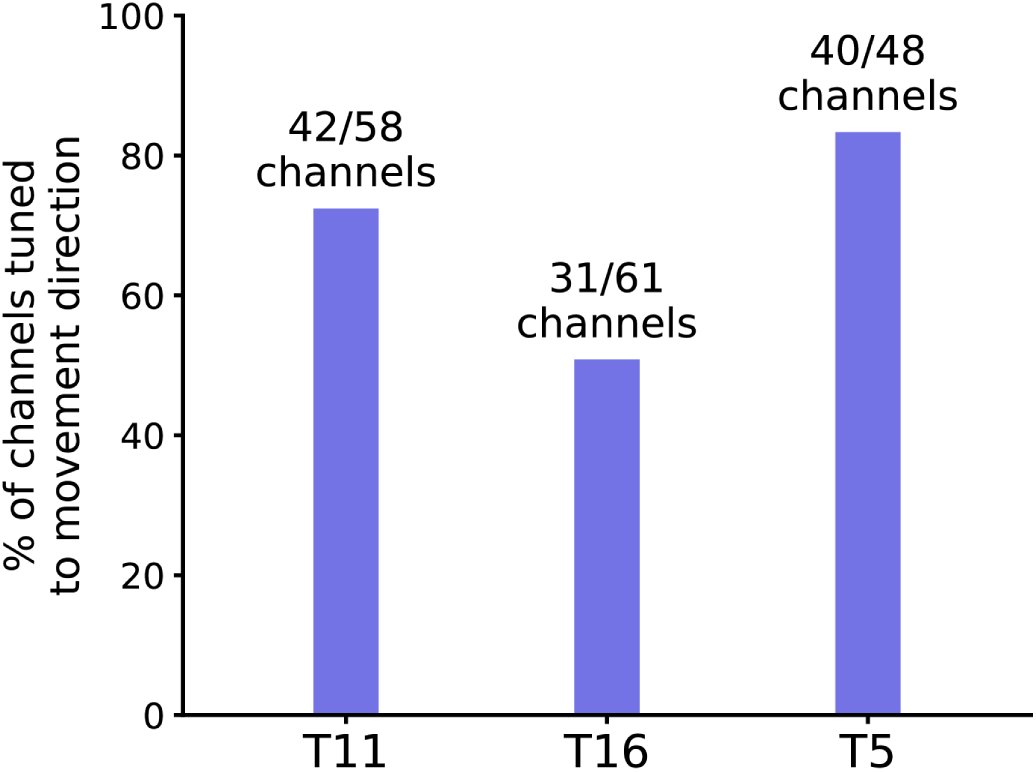
Preparatory tuning to movement direction is prominent across the recorded neural population. Percentage of active channels from participants T11, T16, and T5 with preparatory activity tuned to upcoming movement direction during the radial-8 task (*p*-value < 0.01, ANOVA).

**Supplemental Fig. 2:**
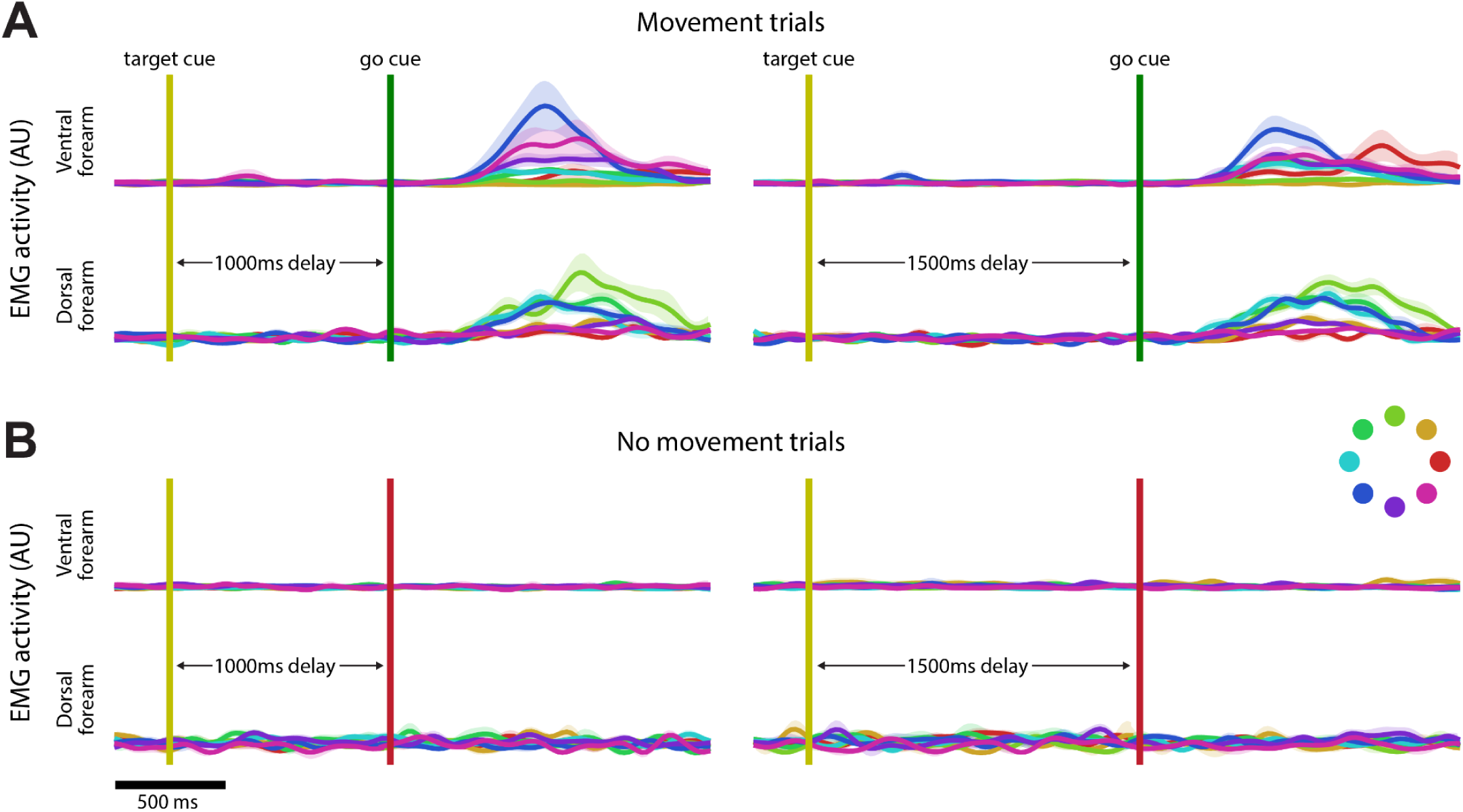
Residual muscle activity is only observed during the cued movement execution period. **A.** Surface electromyography (EMG) recordings from participant T16’s forearm during the instructed-delay radial-8 task for trials where movement was cued following the delay period. Traces correspond to average EMG activity (arbitrary units) through time for trials from the different movement directions (as indicated by the color wheel), and with different delay durations (left and right columns). Shaded regions indicate +/-the standard error of the mean for each condition. Top and bottom rows are surface EMG recordings from T16’s ventral and rostral forearm respectively. **B.** Same as A, but for trials where the movement was skipped.

**Supplemental Fig. 3.**
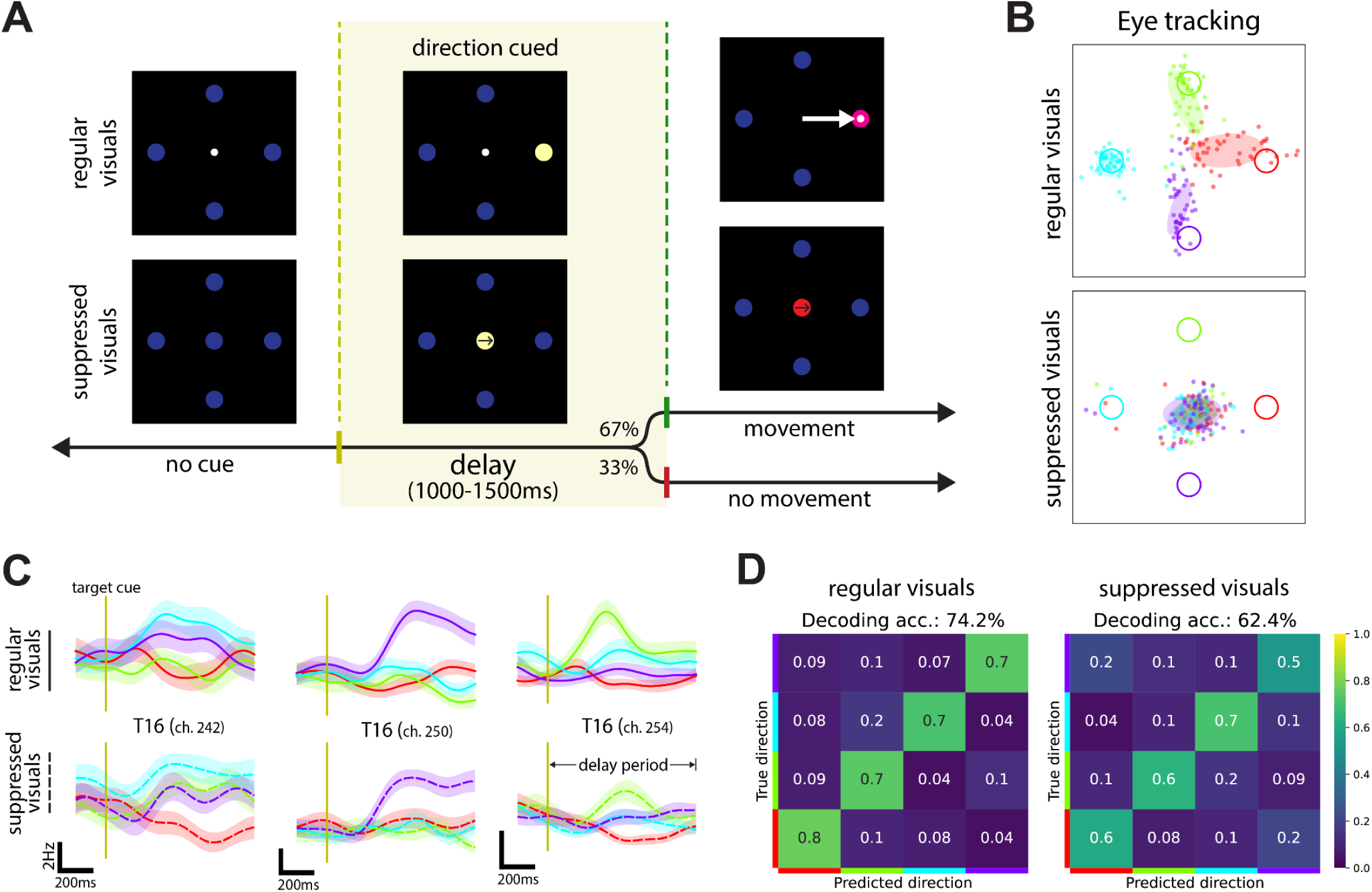
Preparatory activity is still present when target visual saliency and eye movements are non-prominent during the delay period. **A.** Trial structure, timing, and example trial visuals for the regular and alternate versions of the instructed-delay radial-4 tasks designed to test the effect of visual saliency and eye movements on preparatory activity. In the alternate version of the task with suppressed visuals, movement direction was cued by an arrow at the center of the screen, and the participant was instructed to maintain their gaze on the center cue. **B.** Gaze position in screen coordinates recorded using an eye tracker while participant T16 performed both versions of the task. Dots indicate the time-averaged gaze position for each trial (averaged from 600 to 1000 ms after the target cue), colored according to the corresponding cued target direction. Target positions in screen coordinates are indicated by the colored rings. Ellipsoids indicate the mean and +/-one standard deviation of the tracked gaze position across trials for each condition. **C.** Average firing rates through time during the delay period for example neural channels from participant T16, for trials from the task with regular visuals (top row) or suppressed visuals (bottom row). Traces are colored by movement direction (as indicated in B) and shaded regions indicate +/-the standard error of the mean for each condition. **D.** Confusion matrices for single-trial movement direction predictions decoded from preparatory activity for participant T16, using SVM classifiers (leave-one-out cross-validation). SVMs classifiers are fit and evaluated separately for trials from the different tasks (regular and suppressed visuals).

**Supplemental Fig. 4:**
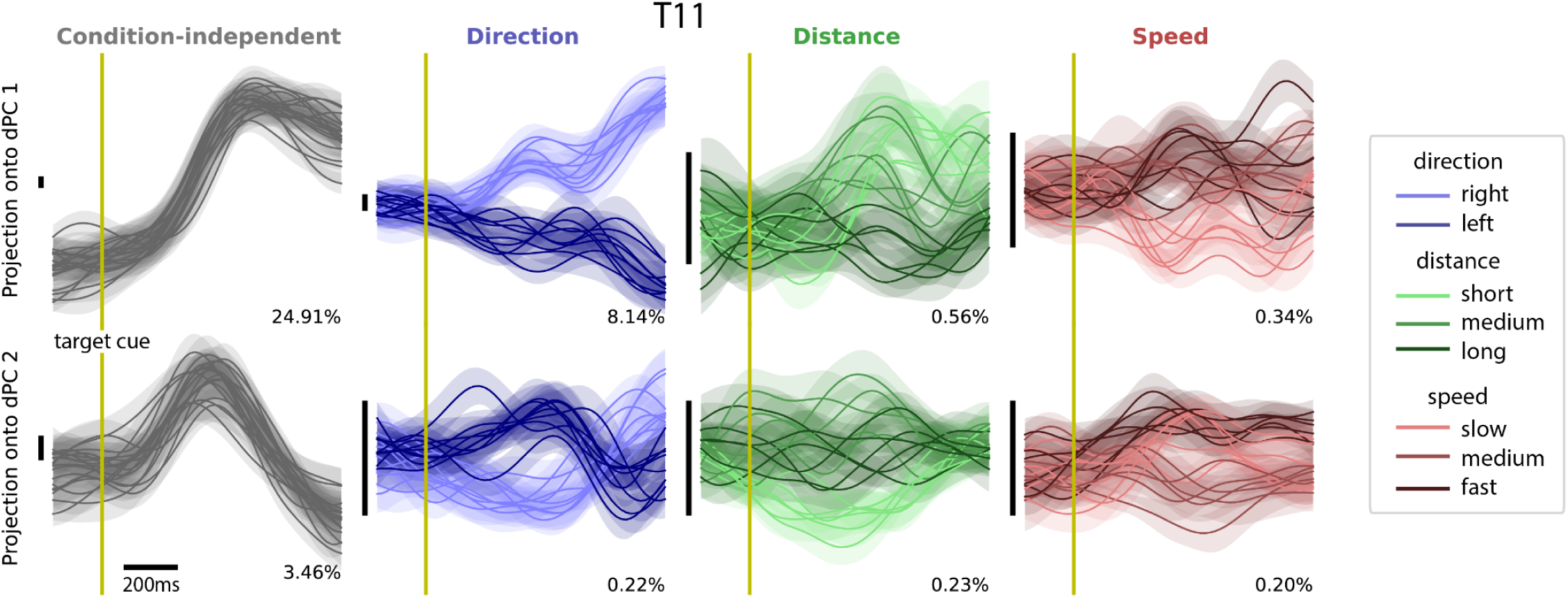
dPCA decomposition of preparatory activity during the instructed-delay Pac-Man task for participant T11. Projection of average neural population activity during the delay period (for participant T11) onto demixed principal components (dPCs) that are condition-independent or specific to upcoming movement direction, distance, or speed. Traces are colored according to the value of the movement feature tied to each specific dPC and shaded regions indicate +/-the standard error of the mean for projections corresponding to trials from each condition. Vertical black bars to the left of each subplot indicate a consistent scale across dPC projections (arbitrary units). The number on the bottom right of each subplot indicates the percentage of total preparatory activity variance explained by each component.

**Supplemental Fig. 5:**
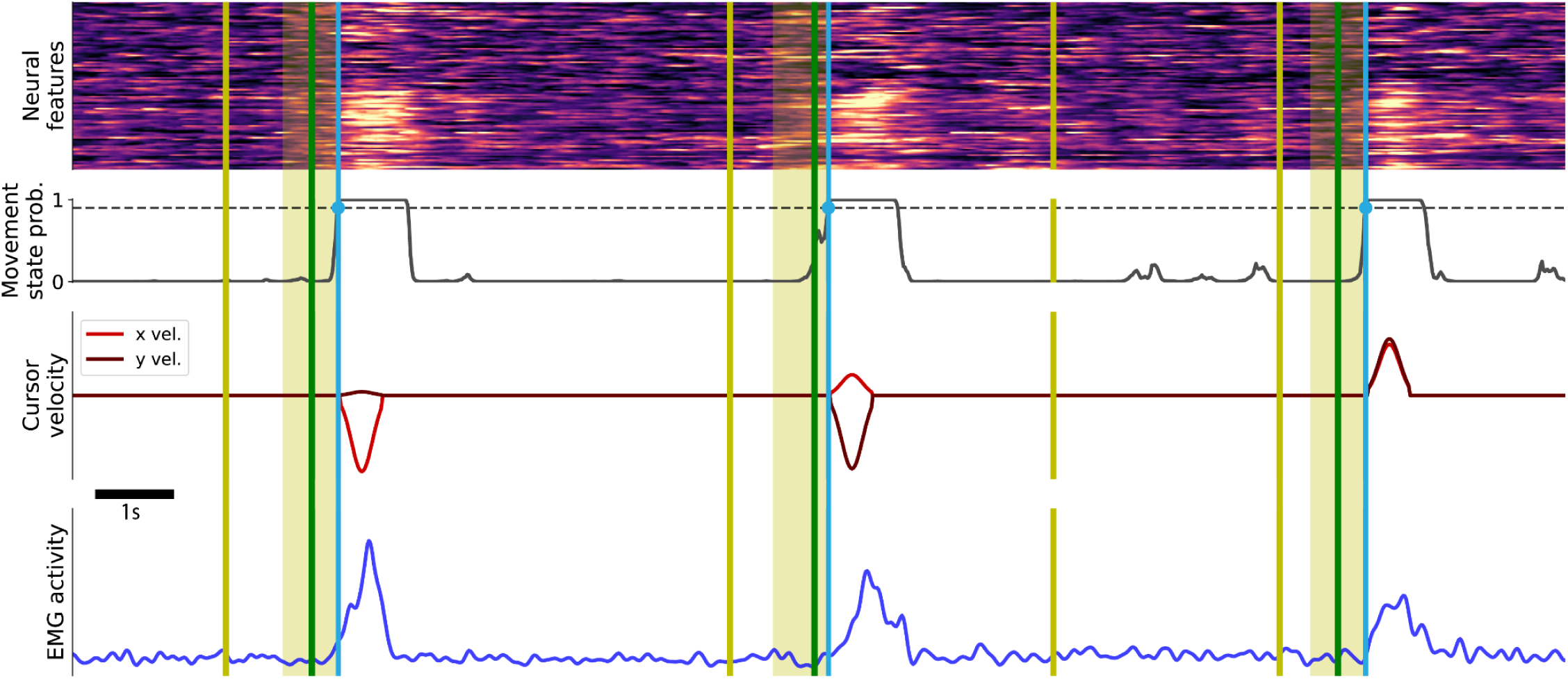
Movement timing inferred by preparatory control paradigm is consistent with recorded EMG activity. First three rows show an example time window with the normalized neural features, decoded movement state probability, and cursor velocity during online evaluation of the preparatory control paradigm with participant T16. Bottom row shows example EMG activity (arbitrary units) recorded from the forearm of participant T16 during the same time window. Yellow lines, green lines, and light blue lines indicate target cue times, go cue times, and decoded movement onset times, respectively. Yellow boxes indicate the windows of preparatory activity used to decode intended movement direction.

